# pyFOOMB: Python Framework for Object Oriented Modelling of Bioprocesses

**DOI:** 10.1101/2020.11.10.376665

**Authors:** Johannes Hemmerich, Niklas Tenhaef, Wolfgang Wiechert, Stephan Noack

**Affiliations:** Institute of Bio- and Geosciences - IBG-1: Biotechnology, Forschungszentrum Jülich GmbH, 52425, Jülich, Germany; Computational Systems Biotechnology (AVT.CSB), RWTH Aachen University, Aachen, 52074, Germany; Bioeconomy Science Center (BioSC), Forschungszentrum Jülich, Jülich, 52425, Germany

**Keywords:** Python, bioprocess modelling, object oriented modelling, ODEs

## Abstract

Quantitative characterization of biotechnological production processes requires the determination of different key performance indicators (KPIs) such as titer, rate and yield. Classically, these KPIs can be derived by combining black-box bioprocess modelling with non-linear regression for model parameter estimation. The presented pyFOOMB package enables a guided and flexible implementation of bioprocess models in the form of ordinary differential equation systems (ODEs). By building on Python as powerful and multi-purpose programming language, ODEs can be formulated in an object-oriented manner, which facilitates their modular design, reusability and extensibility. Once the model is implemented, seamless integration and analysis of the experimental data is supported by various Python packages that are already available. In particular, for the iterative workflow of experimental data generation and subsequent model parameter estimation we employed the concept of replicate model instances, which are linked by common sets of parameters with global or local properties. For the description of multi-stage processes, discontinuities in the right-hand sides of the differential equations are supported via event handling using the freely available assimulo package. Optimization problems can be solved by making use of a parallelized version of the generalized island approach provided by the pygmo package. Furthermore, pyFOOMB in combination with Jupyter notebooks also supports education in bioprocess engineering and the applied learning of Python as scientific programming language. Finally, the applicability and strengths of pyFOOMB will be demonstrated by a comprehensive collection of notebook examples.

## Introduction

Biotechnological production processes leverage the microorganisms’ synthesis capacity to produce complex molecules that are hardly accessible by traditional chemical synthesis. Importantly, modern genetic engineering methods allow for targeted modification of single enzymes and whole metabolic pathways for biochemically accessing value-added compounds beyond those naturally available. However, to render the production of a target compound economically feasible, a suitable bioprocess needs to be developed which fits to an engineered microbial producer strain. In this context, computational modelling approaches utilize existing knowledge on strain and process dynamics, giving rise to modern systems biotechnology. Once a digital representation of a biotechnological system has been implemented, in-silico optimizations can be performed to design an improved bioprocess, effectively reducing the number of wet-lab experiments. With the availability of new experimental data the computational model can be refined to increase its predictive power towards an optimal bioprocess.

Considering the highly interdisciplinary nature of systems biotechnology requiring expertise in (micro-)biology, process engineering, computer science, and mathematics, it becomes obvious that rarely a single person can have a deep knowledge in all these fields. The more specialized and performant a bioprocess model is intended to be, the higher the knowledge level needed by the user. This may prevent non-experts in modeling and programming from dealing with these highly rewarding topics. Consequently, there is a need for tools in systems biotechnology that can be quickly learned and applied by non-experts, with the development of additional skills determined by demand.

Here, we present the pyFOOMB package that enables the implementation of bioprocess models as systems of ordinary differential equations (ODEs) via the multi-purpose programming language Python. Based on the object-oriented paradigm, pyFOOMB provides a variety of classes for the rapid and flexible formulation, validation and application of ODE-based bioprocess models. Table 1 gives a comparative, non-exhaustive overview of software packages that are suitable for bioprocess modelling. These tools were developed with partly other application areas in mind, e.g., modeling and analysis of biochemical networks or simulation of chemical engineering unit operations. Consequently, these software packages require different levels of programming skills and some domain-specific knowledge for accessibility. Therefore, a major driver to establish pyFOOMB was to provide a flexible modelling tool that requires only basic programming knowledge and thus shows low hurdles for beginners in bioprocess modelling. The latter is supported by a comprehensive collection of ready-to-use working examples which come along with pyFOOMB.

**Table 1.**
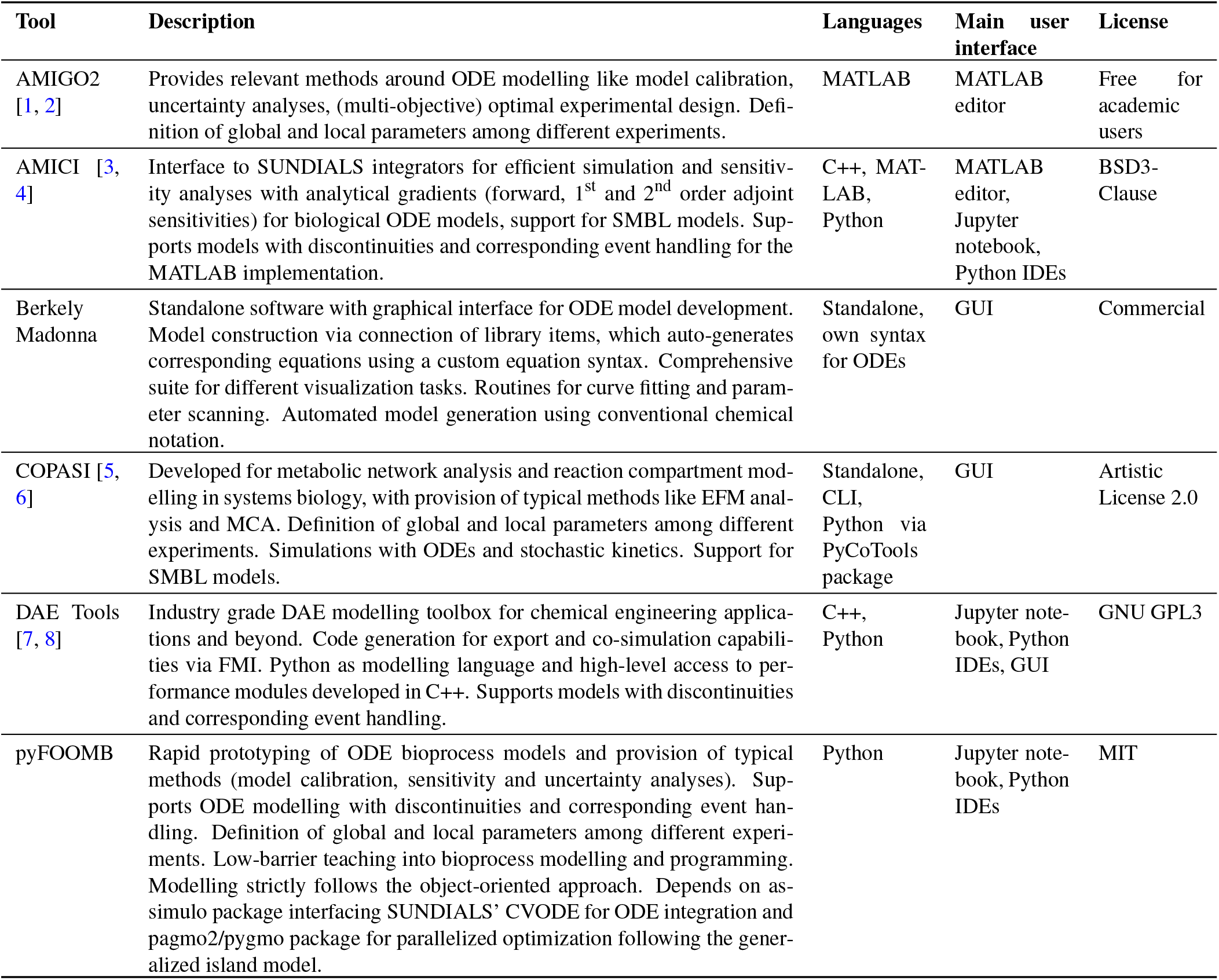
Non-exhaustive comparison of software packages suitable for bioprocess modelling. The listed tools were developed for different application areas and address different primary needs. Therefore, different domain-specific knowledge and programming skills are required for the packages’ accessibility. All packages provide at least several functionalities required for bioprocess modelling.

Due to the full programmatic access to Python, complex models can also be implemented. Furthermore, great importance was given to convenient visualization methods that facilitate the understanding of qualitative and quantitative model behavior. Finally, the enormous popularity of Python as the de-facto standard language for data science applications makes it easy to integrate pyFOOMB with other advanced tools for scientific computing.

## Main functionalities of pyFOOMB for bioprocess modelling

Bioprocess models are implemented as ODEs for the time-dependent variables *x*(*t*):

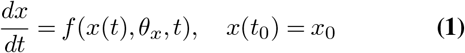

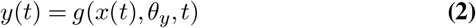

which depend on some model parameters *θ_x_* and initial values *x*_0_. In practice, some of the variables might not be directly measurable. Therefore, observation (or calibration) functions *y*(*t*) can be defined that relate these variables to the observable measurements, thus introducing some additional parameters *θ_y_* into the model.

In order to make the user familiar with our pyFOOMB tool, a continuously growing collection of Jupyter notebook examples is provided. These demonstrate basic functionalities and design principles of pyFOOMB and serve as blueprint for the rapid set up of case-specific bioprocess models (Table A1).

## Modelling workflow when using pyFOOMB

In the following we present a typical workflow for implementing and applying bioprocess models with pyFOOMB (Fig. 1). Throughout this section the toy example model of Figure 2A will be employed.

**Fig. 1.**
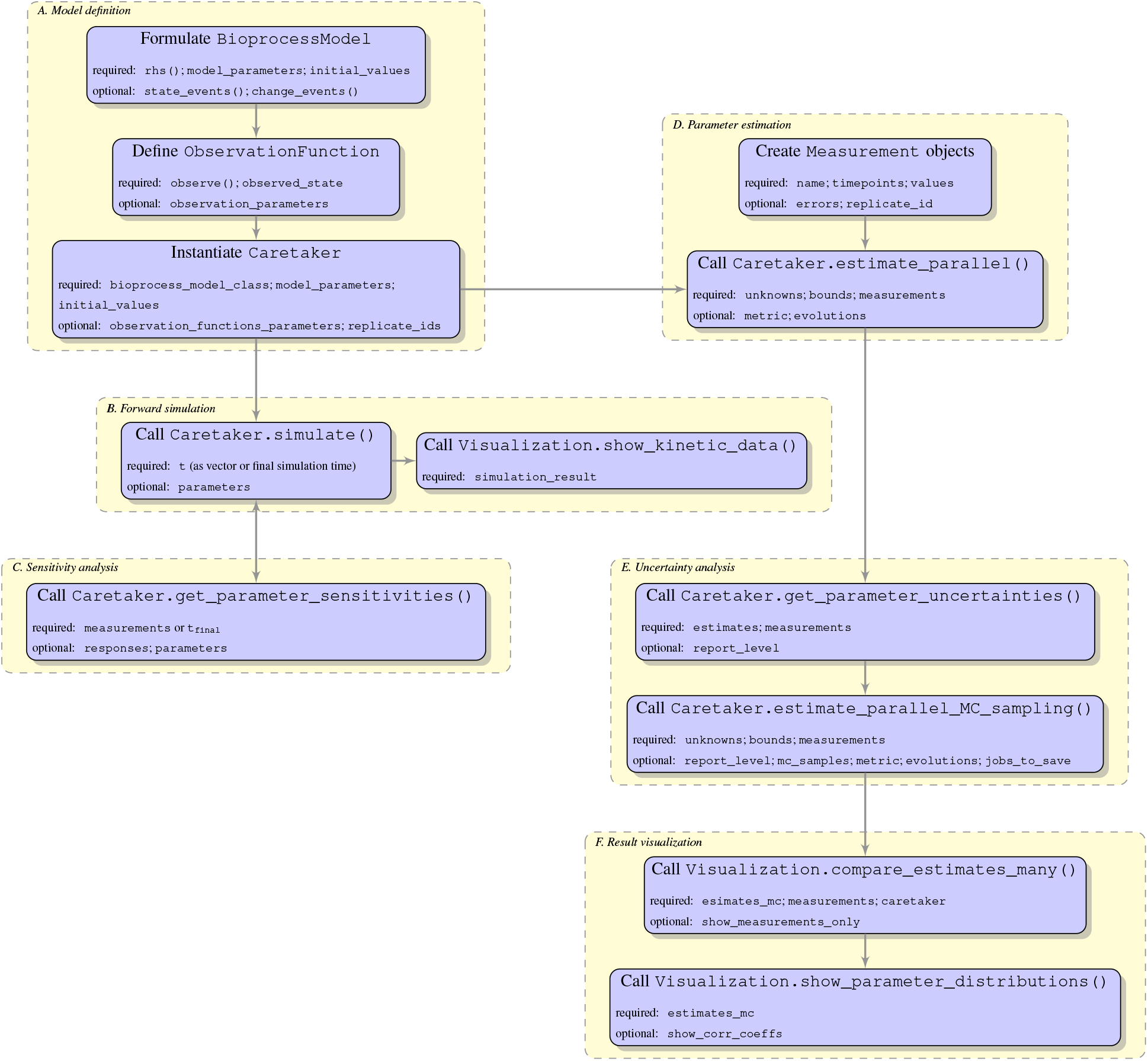
High-level description of a typical bioprocess modelling workflow with pyFOOMB. For a full description of all classes and methods including a complete list of all arguments and default values, please see the provided Jupyter notebook examples and source code documentation.

**Fig. 2.**
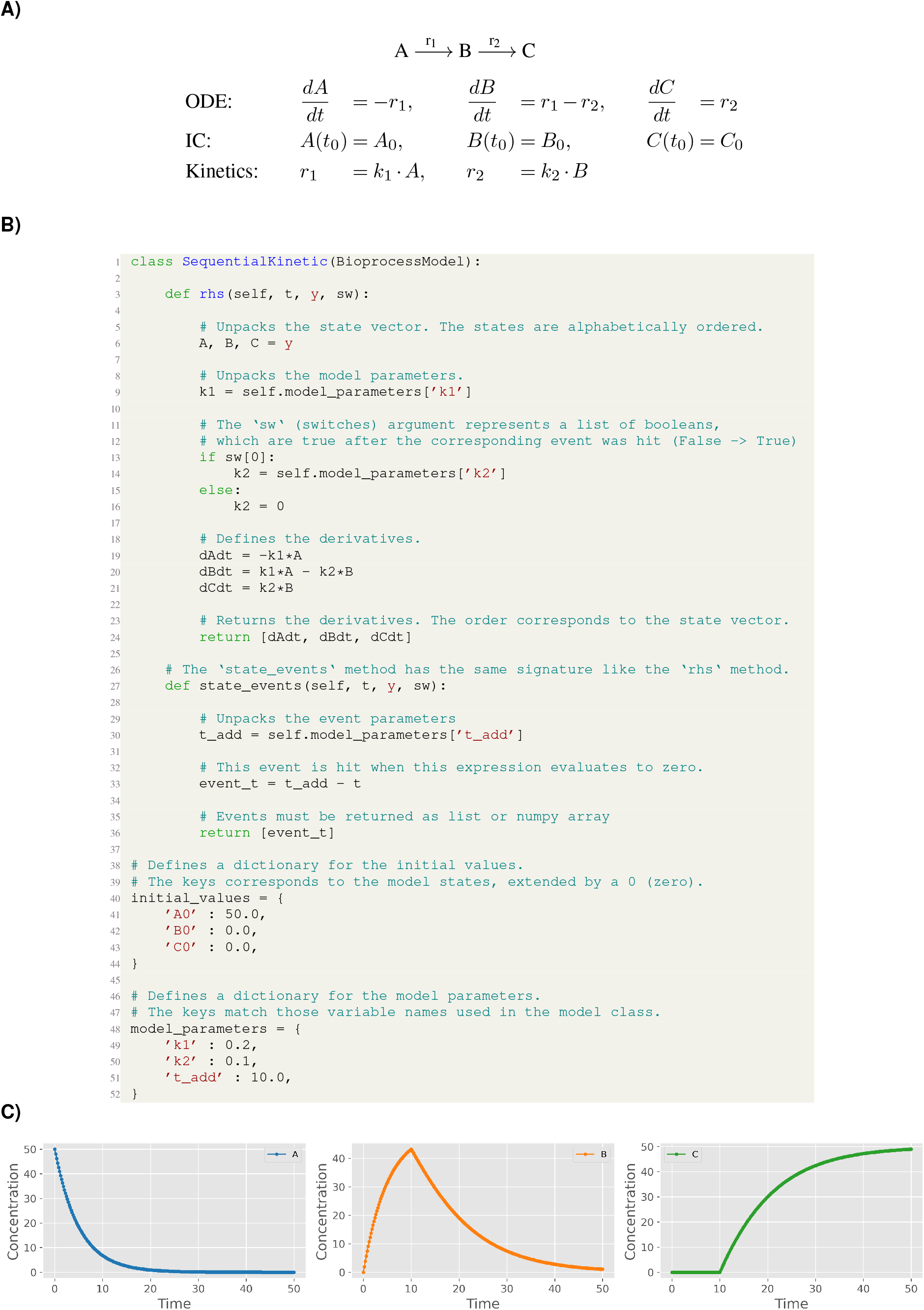
Toy example of a sequential reaction cascade. A) Mathematical representation of the ODE system with initial conditions (IC). B) Object-oriented implementation in pyFOOMB. The ODE is defined within the `rhs`() method. Initial values and model parameters are defined as dictionaries. C) Results of a forward simulation. At *t* = 10 an event occurs, where the conversion from B to C is switched on, i.e. *k*_2_ > 0.

### A. Model definition

In a first step, the targeted model and its parametrization is implemented by creating a user-specific subclass of the provided class `BioprocessModel` (Fig. 2B). This basic class provides all necessary methods and properties to run forward simulations for the implemented model. Essentially, the abstract method `rhs`() must be formulated by the user.

#### Discrete behavior

To monitor and control the dynamics of specific model variables so called `state_events`() and `change_states`() methods can be defined. This is for example required for the modelling of multi-phased processes such as fed-batch with event-based changes in feeding regimes.

#### Observation of model states

In order to connect the model variables to measurable quantities, an `ObservationFunction` can be created, with the mandatory implementation of the `observe`() method for each relevant calibration function. Noteworthy, a variable’s state can be linked to different observation functions, reflecting the fact that there are typically several analytical methods available for one specific bioprocess quantity. This approach allows to separate the bioprocess model from corresponding observations functions and thus, increases re-usability of the different parts. By deriving initial guesses for the parameters, a forward simulation from the model is typically used to verify the intended qualitative behavior in comparison to the experimental data.

#### Global and local parameters

A key feature of pyFOOMB is the possibility to integrate measurement data from independent experimental runs (replicates) by creating a corresponding number of new instances of the same model. These can still share a common set of model parameters that are defined as “global”, but at the same time differ in some other “locally” defined parameters.

Typical global parameters of an ODE-based bioprocess model are the maximum specific growth rate *μ_max_* or the substrate specific biomass yield *Y*_X/S_, while all initial values are reasonable defined as local parameters (see Application example II). Different values for the local parameters reflect biological or experimental variability that may arise from slight deviations in preparing, running or analyzing each replicate experiment. Alternatively, such variability might be introduced by purpose when conducting replicate experiments with intentionally very different starting conditions. The latter refers to a classical design-of-experiment approach aiming for experimental data with a maximum information gain with respect to the global parameters.

#### Working with the model

The implemented model (including an initial parametrization) is passed to the instantiation of the `Caretaker` class (Fig. 1). During the instantiation procedure several sanity checks run in the back and, in case of failure, direct the user to erroneous or missing parts of the model. The resulting object exposes important and convenient methods typically applied for a bioprocess model, such as running forward simulations, setting parameter values, calculating sensitivities, estimating parameters, and managing replicates of model instances.

### B. Forward simulation

For a certain set of model parameters the time-dependent dynamics of the model variables and corresponding observations are obtained by running a forward simulation (cf. Fig. 1). Integration of the ODE system is delegated to the well-known Sundials CVode integrator with event detection [9]. Its Python interface is provided by the assimulo package [10], which implements seamless event handling hidden from the user. Running some forward simulations with subsequent visualization is a convenient approach to verify the qualitative and quantitative behavior of the implemented model (Fig. 2C). pyFOOMB provides a class with convenient methods for that purpose, e.g, plotting of time series data covering model simulations and measurement data, corner plots for one-by-one comparison of (non-linear) correlations between parameters from Monte-Carlo sampling as well as visualization of the results from sensitivity analysis.

### C. Sensitivity analysis

Local sensitivities *∂_yi_*(*t*)/*∂θ_j_* are available for any model response *y_i_* (model state or observation) with respect to any model parameter *θ_j_* (including ICs and observation functions). The sensitivities are approximated by the central difference quotient using a perturbation value of *h* · *max*(1,|*θ_j_*|). Sensitivities can also be calculated for an event parameter that defines implicitly or explicitly a point in time where the behaviour of the equation system is changed (cf. Fig. 3A). This is useful for, e.g., analyzing induction profiles of gene expression or irregular bolus additions of nutrients.

**Fig. 3.**
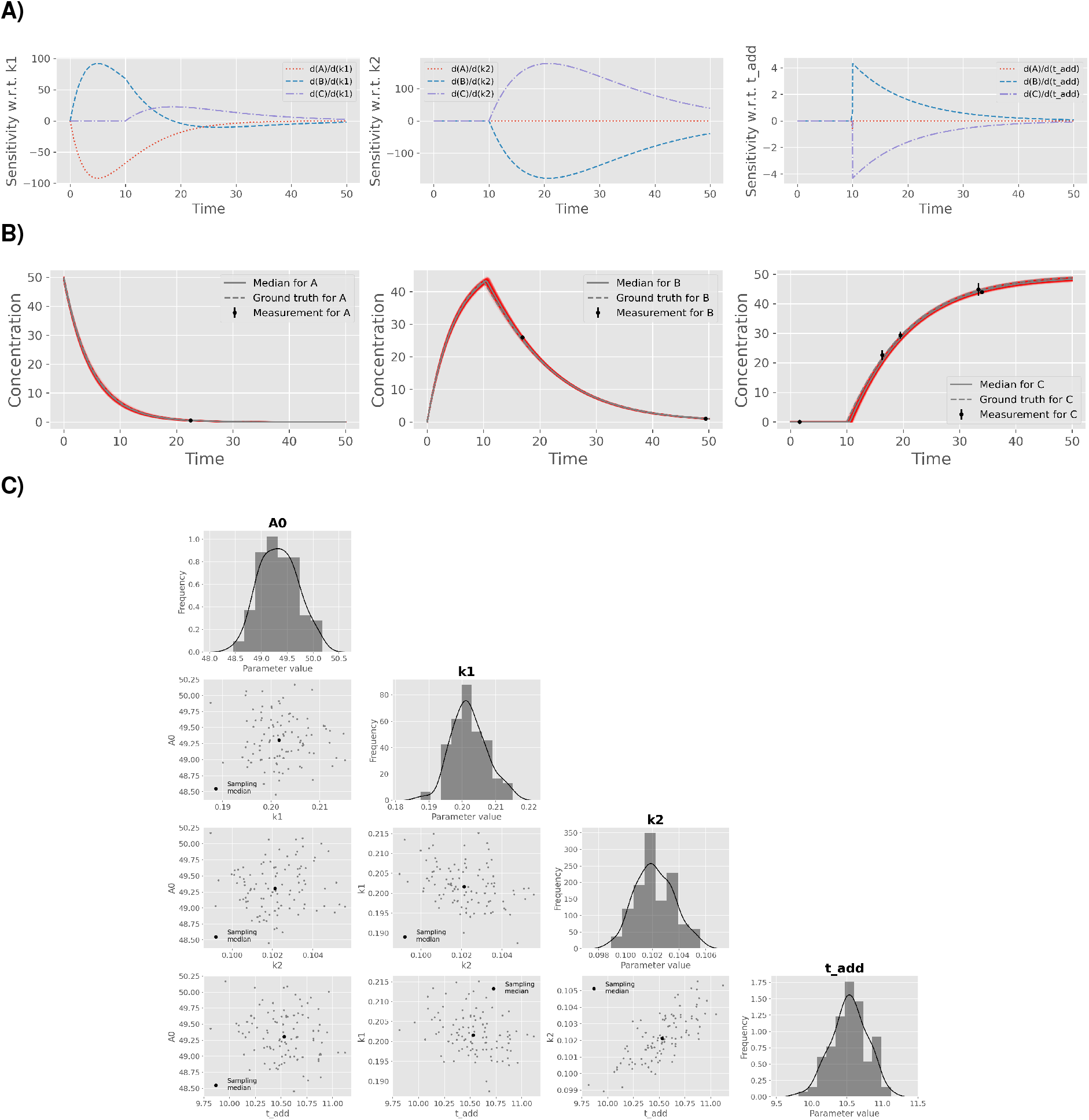
Essential steps of model validation supported by pyFOOMB. A) Sensitivity analysis of the model states with respect to the three parameters *k*_1_, *k*_2_ and *t*_add_. B) Parameter estimation using artificial experimental data with random noise (black dots with error bars) in combination with parallelized MC sampling (red lines). The median of 125 single parameter estimations is shown in grey. C) Uncertainty analysis using a corner plot of the resulting empirical parameter distributions. Diagonal elements show the individual distributions as histogram with a kernel density estimate, while off-diagonal elements indicate one-by-one comparisons of each parameter pair. The plot was generated using the `show_parameter_distributions`() method of pyFOOMB’s `Visualization class`.

### D. Parameter estimation

Finding those parameter values for a model that describe a given measurement dataset best is implemented as a typical optimization problem. Here, the `estimate_parallel`() method is the first choice, because it employs performant state-of-the-art meta-heuristics for global optimization, which are provided by the pygmo package [11]. In contrast to local optimization algorithms, there are no dedicated initial guesses needed for the parameters to be estimated (“unknowns”). Instead, lower and upper estimation bounds are required. As a good starting point such bounds can be derived from explorative data analysis (see Application example II), literature research, or expert knowledge by simply assuming three orders of magnitude centered around the precalculated or reported parameter value.

Noteworthy, pygmo provides Python bindings to the pagmo2 package written in C++. It implements the asynchronous generalized island model [12], which allows to run several, different algorithms cooperatively on the given parameter estimation problem. As an inherent feature of this method, an optimization run can be executed for a given number of so called “evolutions” and after inspection of the results, the optimization can be continued from the best solution found so far (Fig. 3B). This powerful approach allows to traverse multi-modal, non-convex optimization landscapes.

Currently, the maximum likelihood estimators (covering its classical variants least-squares and weighted-least-squares) are implemented. In general, a parameter vector 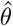 is to be found that minimizes a certain optimization (loss) function. For example, for the negative log-likelihood (NLL) function for normally distributed measurement errors it holds:

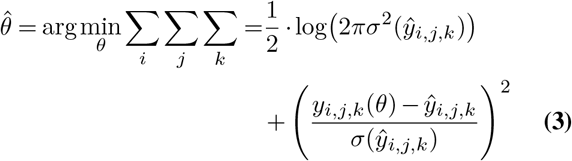

Given a specific measurement 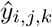, for each corresponding model response *i* at sampling time point *j* and replicate *k*, the NLL is calculated and summed up. By default, it is assumed that all measurements follow normal distributions based on mean values and corresponding standard deviations. The loglikelihood function is constructed by pyFOOMB when starting the parameter estimation procedure. For the case that measurements are assumed to follow other distributions, this can be specified when creating the `Measurement` object and pyFOOMB will take care for the definition of the correct log-likelihood function.

Noteworthy, it is not required to provide complete measurement datasets, i.e. a specific replicate may contain only one measurement or even unequal data points for different model responses.

### E. Uncertainty analysis

An approximation of the parameters’ variance-covariance matrix is provided by inversion of the Fisher information matrix, which is calculated from local sensitivities (see above). Besides, non-linear error propagation is available by running a repeated parameter estimation procedure starting from different Monte-Carlo samples (so called “parametric bootstrapping”, Fig. 3C). A parallelized version of this method is provided based on the pygmo package.

### F. Result visualization

Following parameter estimation and uncertainty analysis via parametric bootstrapping, (non-)linear correlations between each pair of parameters can be readily visualized with the method `show_parameter_distributions`(). In addition, results are typically inspected by visualizing the set of model predictions according to the calculated parameter distributions. Using the `compare_estimates_many`() method, a direct comparison between measurements and repeated simulations is possible, which makes it easier to assess the validity of the model.

### G. Implementation of model variants

Usually, when starting to formulate a bioprocess model there is not only one option to link a specific rate term with a suitable kinetic model. Depending on how informative the available measurements are in relation to the unknown kinetics, it could make sense to directly start the whole workflow by setting up a “model family”.

Following the object-oriented approach of pyFOOMB, a model family can be easily set up based on inheritance (Fig. 4A). In principle, for each relevant part of the original model additional submodels can be introduced by declaring separate methods. In a programming context, this approach is also known as “method extraction”, as the calculations in question are extracted into further dedicated methods. The model family is then realized by building on a common model structure encoded in the `BaseModel` and a set of subclasses encoding the specific submodels. On a technical level, the definition of “abstract” methods is required to enforce the individual members of the model family to implement their specific submodel.

**Fig. 4.**
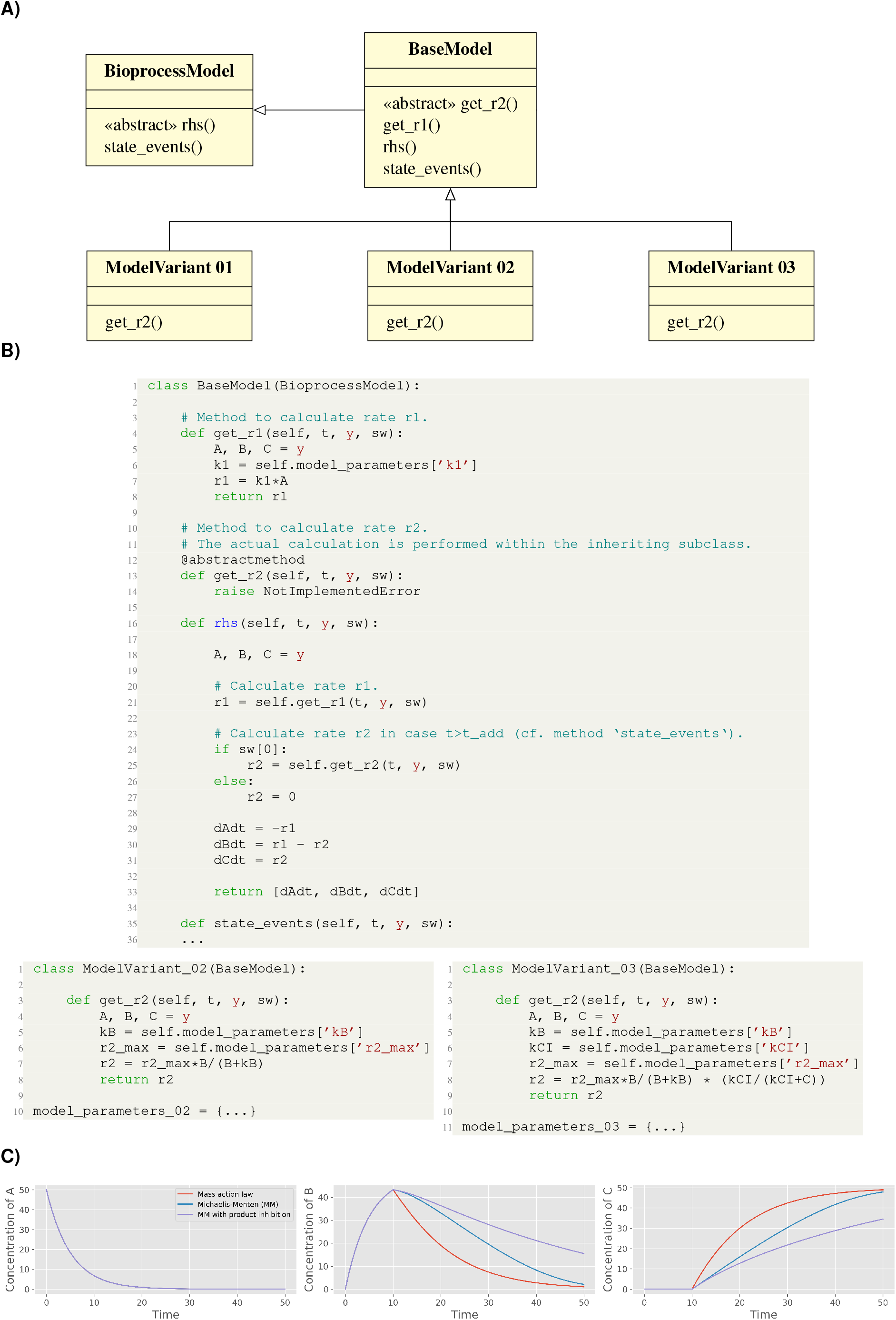
Implementation of model variants using inheritance. A) UML class diagram for three model variants of the toy model. The kinetic rate law for reaction *r*_2_ is set as either Mass action, Michaelis-Menten, or Michalis-Menten with product inhibition. B) Python implementation of the base class `BaseModel` with the abstract method `get_r2`() and two example subclasses. (C) Resulting forward simulations comparing the model variants.

In an extended version of the running example, the rhs() method of the `BaseModel` class now depends on the two additional methods `get_r1`() and `get_r2`() to separate the calculation of rates *r*_1_ and *r*_2_, respectively (Fig. 4B). The latter is declared as an abstract method to enable a family of models (`ModelVariant01-03`) for comparing different rate expressions of *r*_2_.

In the following sections two different applications examples will be presented that apply the introduced modelling workflow of pyFOOMB.

## Application example I: Small-scale repetitive batch operation

In the first example workflow specific growth rates within an Adaptive Laboratory Evolution (ALE) process are determined. ALE processes utilize the natural ability of microorganisms to adapt to new environments to improve certain strain characteristics, such as growth on a specific carbon source.

Here, a *Corynebacterium glutamicum* strain which was able to slowly (*μ*_max_ < 0.10 h^-1^) utilize D-xylose, was cultivated repeatedly in defined medium containing D-xylose as sole carbon and energy source. The cultivation was done in an automated and miniaturized manner, delivering a biomass-related optical signal, “backscatter”, with a high temporal resolution. This signal was used to automatically start a new batch from the previous one, as soon as a backscatter threshold was reached. The threshold was deliberately choosen to be in the mid-exponential phase, where no substrate limitation was to be expected. Six individual clones were cultivated over one precculture and seven repetitive batches, as shown in Fig. 5A.

**Fig. 5.**
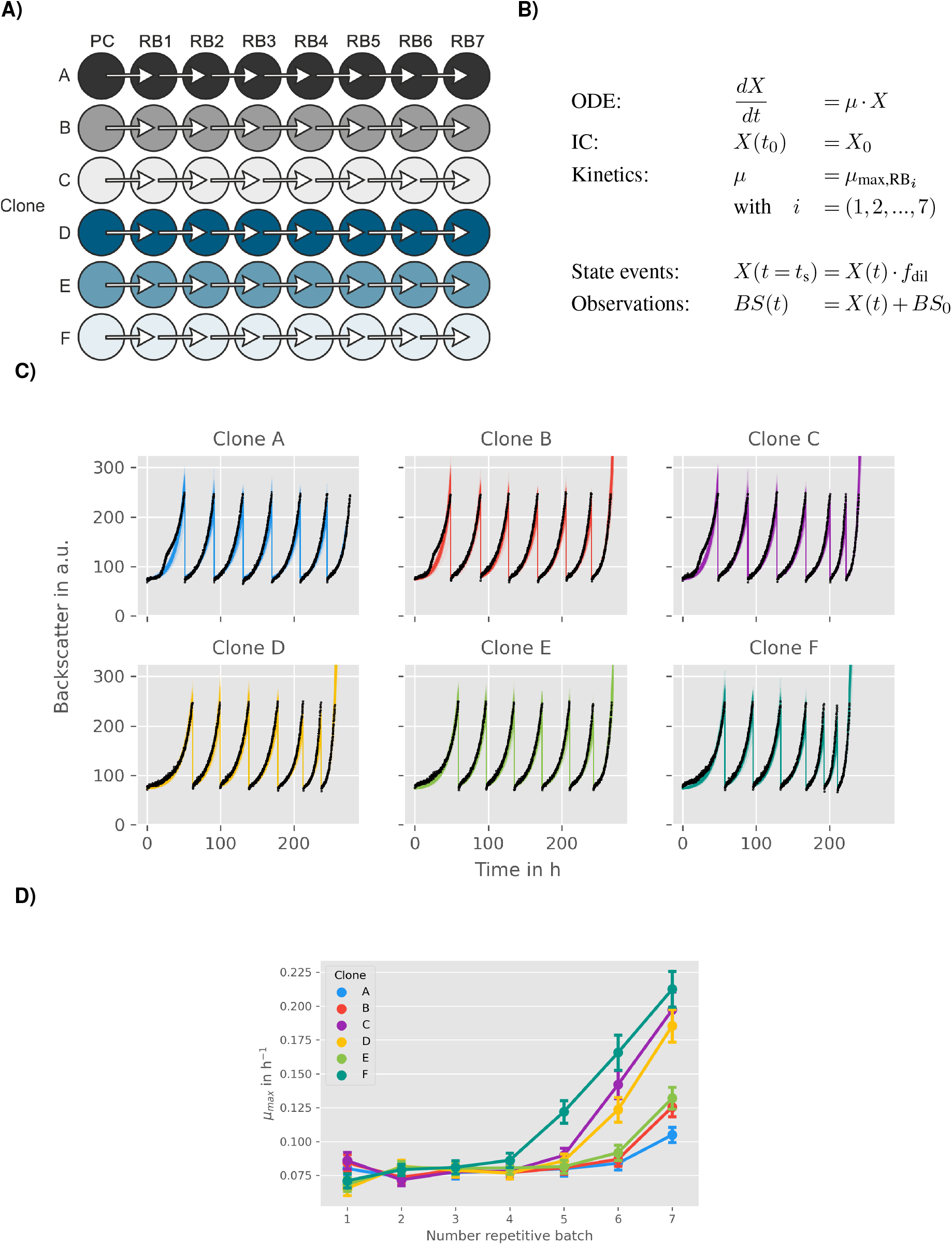
Modelling and analysis of small-scale repetitive batch processes. A) Experimental layout for fully automated repetitive batch operation in microtiter plates (taken from [13]. Each cycle was started from 6 independent clones followed by 7 consecutive batches. B) ODE model for describing the biomass dynamics including state events for multiple sampling and growth rate estimation. C) Time course of online backscatter data (black dots) and corresponding model fits (straight coloured lines). D) Evolution of maximum specific growth rates in each cycle. Mean values and standard deviations were estimated by parallelized MC sampling (*n* = 200).

### Model development

In order to keep the number of parameters and computation times as low as possible, a rather simple bioprocess model as shown in Fig. 5B was employed.

Growth is determined solely by the growth rate *μ*. Substrate limitations are not taken into account, since the experimental design (see above) should avoid these sufficiently. Biomass *X* is not measured directly, instead, backscatter is introduced to the model via an `ObservationFunction`. This function describes a linear relationship between backscatter and biomass and takes the blank value *BS_0_* of the signal into account. A relative measurement error for the backscatter signal of 5 % is assumed based on expert knowledge. The model describes the whole ALE process for each clone, not an individual batch. Therefore, state events are used to trigger a state change of *X*, where *X* is multiplied by a dilution factor *f*_dil_. Additionally, the maximum growth rate parameter is switched for each repetitive batch. As a result, an individual *μ*_max_ for each repetitive batch and each clone is gained. Since initial inoculation of the different clones and the inoculation procedure within the experiment was the same for all, initial biomass concentration *X*_0_ and dilution *f*_dil_ are considered as global parameters.

### Parameter estimation and uncertainty analysis

In total, model parameters for six clones are estimated, which form six replicates in the context of pyFOOMBs modelling structure. For each clone, seven maximum growth rates are to be determined, plus *X*_0_, *f*_dil_, and *BS*_0_ as global parameters, thus 44 parameters in total. Parallelized MC sampling was used to obtain distributions for all parameters. Results are shown in Fig. 5C and D.

The estimated backscatter signals follow the actual data closely, resulting in narrow distributions for the parameters of interest, the individual *μ*_max_ values for each clone and repetitive batch. For example, clone F starts with growth rates of 0.071 ± 0.005 h^-1^ to 0.086 ± 0.005 h^-1^ for the first four batches. In the fifth batch, a notable raise in maximum growth rate to 0.122 ± 0.008 h^-1^ is visible, indicating one or more beneficial mutation events. Finally, clone F reaches a growth rate of 0.212 ± 0.013 h^-1^. Overall, the estimated growth rates are in good agreement with findings from the original paper.

In another style of ALE experiment, which is not subject in this study, a subpopulation of cells with beneficial mutations was enriched, yielding strain WMB2_evo_, which is analyzed in the second application example.

## Application example II: Lab-scale parallel batch operation

In this example workflow some KPIs of an engineered microbial strain cultivated in a bioreactor under batch operation are determined. Often, such KPIs represent process quantities that are not directly measurable (e.g., specific rates for substrate uptake, biomass and product formation) and therefore have to be estimated using a model-based approach.

The data originates from two independent cultivation experiments with the evolved *C. glutamicum* strain WMB2_evo_ as introduced before [13]. Following successful adaptive laboratory evolution this strain has now improved properties for utilizing D-xylose as sole carbon and energy source for biomass growth. At the same time the strain produces significant amounts of D-xylonate, a direct oxidation product of D-xylose.

### Explorative data analysis and model development

Before implementing a suitable bioprocess model with pyFOOMB, the data from one replicate bioreactor cultivation is visualized and used for explorative data analysis. In Figure 6A the time courses of biomass (*X*), D-xylose (*S*), and D-xylonate (*P*) are presented in one subplot. It can be seen that biomass formation stops with depletion of D-xylose and, thus, modelling the cell population growth by a classical Monod kinetic is reasonable (Fig. 6B). The formation of D-xylonate is also strictly growth-coupled, leading to a simple rate equation with the yield coefficient *Y_P/X_* as proportionality factor. Finally, the D-xylose uptake rate equals the combined carbon fluxes into biomass and D-xylonate, which are related to the yield coefficients *Y_X/S_* and *Y_P/S_* respectively.

**Fig. 6.**
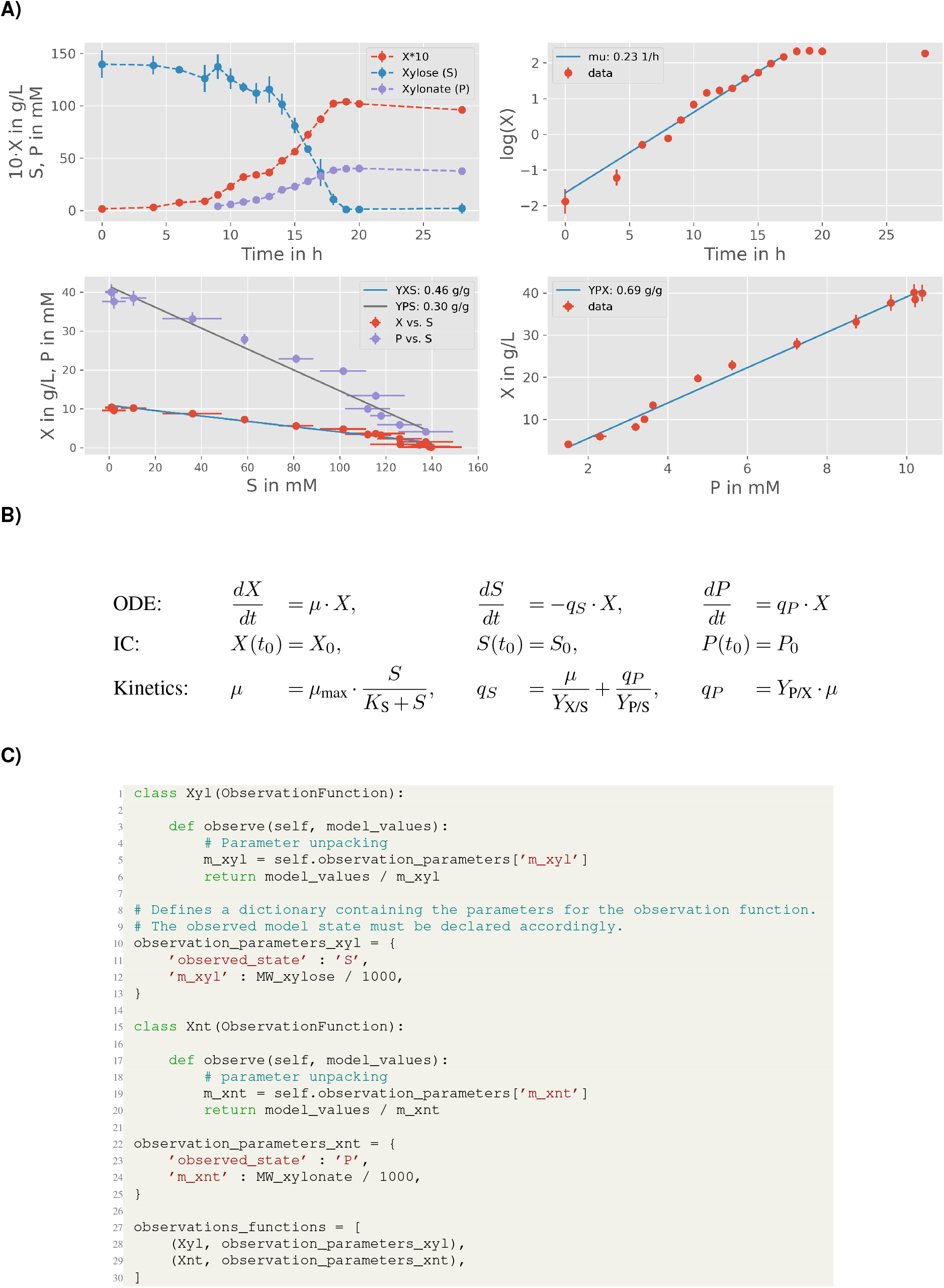
Modelling of lab-scale batch processes. A) Explorative data analysis for one replicate culture. Concentrations for biomass, D-xylose and D-xylonate are denoted by symbols *X*, *S* and *P*, respectively. Following linear regression analysis first estimates for the model parameters *Y*_x/s_, *Y*_p/s_ and *Y*_p/x_ can be derived (for later comparison values are transformed to mass-based units). B) ODE model using classical rate equations. C) Formulation of specific observation functions to map the state variables to the measurements. Here simple transformations from measured molar concentrations to simulated mass concentrations are performed.

The time courses of substrate and product are measured in molar concentrations, while the bioprocess model is formulated using mass concentrations of the respective species. The mappings are realized by defining corresponding observation functions (Fig. 6C).

Finally, the strain-specific parameters like *μ*_max_ and *Y_X/S_* are defined as global parameters, while experiment-specific pa-rameters (ICs for biomass *X* and substrate *S*) are defined as local parameters since the cultivation media and inoculation material were prepared individually for each reactor. Please note, even this very simple process model now already contains eight model parameters (i.e., three ICs and five kinetic parameters) that have to be estimated from the given measurements.

### Parameter estimation and uncertainty analysis

In order to facilitate the parameter estimation problem, good initial guesses for all parameter values are important. First approximations for *μ*_max_ as well as all yield coefficients can be derived by following ordinary and orthogonal distance regression analysis on the raw data assuming linear relationships (Fig. 6A). For Python, corresponding methods are available from the NumPy [14] and SciPy [15] packages.

From the obtained initial guesses corresponding parameter bounds are fixed to run a parallel parameter estimation procedure (Fig. 7A). As a result, a first set of best-fitting parameter values is obtained from which new bounds can be derived for the subsequent uncertainty analysis using again parallelized MC sampling. Corresponding results are summarized in Table 2.

**Fig. 7.**
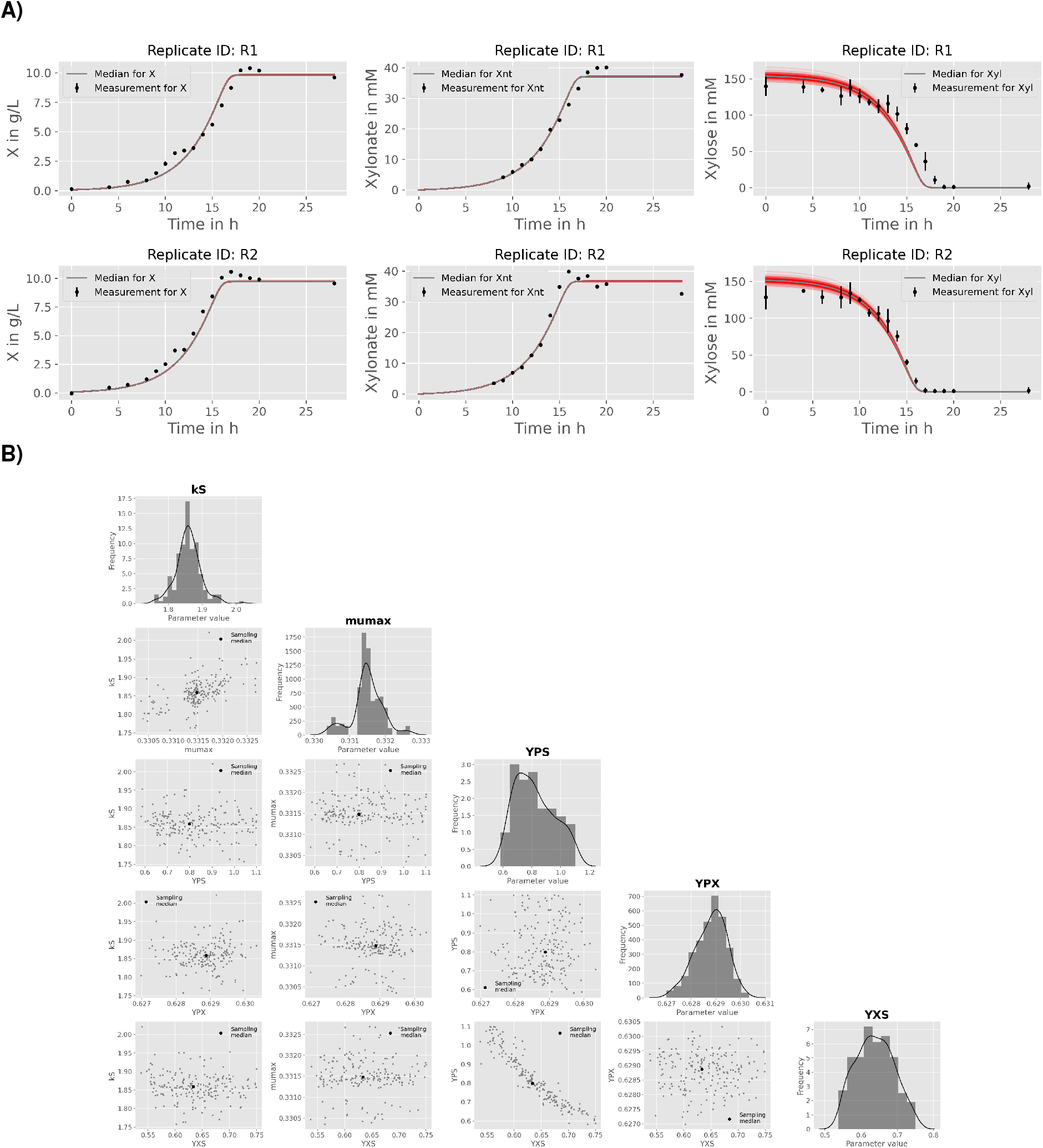
Results from repeated parameter estimation using parallelized MC sampling (*n* = 200). A) Comparison of model predictions with experimental data. B) Uncertainty analysis using a corner plot of the resulting empirical parameter distributions. For the sake of brevity, only the global model parameters are shown.

**Table 2.** Estimated parameter values of the bioprocess model applying parallelized MC sampling. Indices R1 and R2 for parameters *S*_0_ and *x*_0_ indicate their local property following the integration of the two independent replicate experiments.

| Parameter | Property | Unit | Median (16, 84 percentile) |
| --- | --- | --- | --- |
| $k_S$ | global | $g_S L^{-1}$ | 1.86 (1.83 - 1.89) |
| $\mu_{\max}$ | global | $h^{-1}$ | 0.33 (0.33 - 0.33) |
| $Y_{P/S}$ | global | $g_P g_S^{-1}$ | 0.80 (0.68 - 0.99) |
| $Y_{P/X}$ | global | $g_P g_X^{-1}$ | 0.63 (0.63 - 0.63) |
| $Y_{X/S}$ | global | $g_X g_S^{-1}$ | 0.63 (0.58 - 0.69) |
| $S_{0,R1}$ | local | $g_S L^{-1}$ | 23.04 (22.76 - 23.36) |
| $S_{0,R2}$ | local | $g_S L^{-1}$ | 22.78 (22.50 - 23.09) |
| $X_{0,R1}$ | local | $g_X L^{-1}$ | 0.070 (0.070 - 0.071) |
| $X_{0,R2}$ | local | $g_X L^{-1}$ | 0.088 (0.088 - 0.088) |

The pair-wise comparison of parameter distributions shown in 7B reveals a distinct non-linear correlation between the yield coefficients *Y*_p/S_ and *Y*_X/S_. This effect is expected due to the formulation of the biomass-specific substrate consumption rate *q_S_* (Fig. 6B). Equal values for *q_S_* can be derived for different combination of substrate conversion rates into biomass and product, and the yield coefficients are the corresponding scaling factors. The latter is also the reason why the estimated yield coefficients are significantly higher as compared to the explorative data analysis, which does not allow this separation and therefore leads to false-to-low predictions (Table 2 and Fig. 6A).

Finally, the estimated biomass yield *Y_X/S_* for D-xylose is close to the value reported for the wild-type strain growing on D-glucose, i.e. 0.63 [CI: 0.58 - 0.69] vs 0.60 ± 0.04 g_X_ g_S_^-1^ [16]. This indicates a comparable efficiency of **C.** *glutamicum* WMB2_evo_ in utilizing D-xylose for biomass growth.

### Conclusions

The pyFOOMB package provides straight-forward access to the formulation of bioprocess models in a programmatic and object-oriented manner. Based on the powerful, yet beginner-friendly Python programming language, the package addresses a wide range of users to implement models with growing complexity. For example, by employing event methods, pyFOOMB supports the modelling of discrete behaviors in process quantities, which is an important feature for the simulation and optimization of fed-batch processes. The concept of model replicates and definition of local and global parameters mirrors the iterative nature of data generation from cycles of experiment design, execution and evaluation. Moreover, seamless integration with existing and future Python packages for scientific computing is greatly facilitated.

In summary, pyFOOMB is an ideal tool for model-based integration and analysis of data from classical lab-scale experiments to state-of-the-art high-throughput bioprocess screening approaches.

## Availability

The source code for the pyFOOMB package is freely available at github.com/MicroPhen/pyFOOMB. It is published under the MIT license. Currently, its compatibility is tested with Python 3.7 and 3.8, for Ubuntu and Windows operating systems. The use of pyFOOMB within a conda environment is recommended, since the most recent versions of important dependencies are maintained at the conda-forge channel.

## Conflict of interest

The authors have no conflict of interest to declare.

## ACKNOWLEDGEMENTS

This work was partly funded by the German Federal Ministry of Education and Research (BMBF, projects: “Digitalization in Industrial Biotechnology”, grant no. 031B0463 and “BioökonomieREVIER_INNO: Entwicklung der Modellregion BioökonomieREVIER Rheinland”, grant no. 031B0918A). Further funding was received from the Bioeconomy Science Center (BioSC, Focus FUND project “HyIm-PAct - Hybrid processes for important precursor and active pharmaceutical ingredients”, grant no. 313/323-40ø-ø0213).

## Appendix: Notebook examples

**Table A1.** Jupyter notebook examples provided with the pyFOOMB package.

| # | Title | Topic |
| --- | --- | --- |
| 1 | Modelling | Demonstrates basic usage of the pyFOOMB package based on a simple toy mass-action kinetic, i.e. how to implement the right-hand-side of the resulting equation system and how to run and visualize forward simulations. The concept of model replicates is introduced and resulting effects are visualized. |
| 2 | Events | The previous model is extended by events, which are timepoints where the model behavior can be changed safely without interfering with the multistep logic of the ODE integrator. Shows how to use different parameter values before and after an event, as well as how to manipulate the model state values upon reaching an event. |
| 3 | Observation functions | Introduces observation functions that map a model state to an observation, according to a specific, parametrized function. |
| 4 | Parameter estimation | Describes one of the major functionalities of the pyFOOMB package. Parameter values of an implemented model are estimated based on (artificial) noisy data. The presented methods use algorithms from Scipy's optimize module. Besides approximation of uncertainties for estimated parameters based on Fisher information matrix, Monte-Carlo sampling is introduced as method for non-linear error propagation. Suitable visualization methods are used for interpretation of results. |
| 5 | Sensitivities | Shows how to calculate local sensitivities with subsequent visualization. |
| 6 | Bioprocess models | Implements several example bioprocess models, serving as starting point for implementation of user-specific ones. |
| 7 | Fed-batch models | Demonstrates the implementation of fed-batch bioprocess models at various complexities. Shows how to use the models to derive further performance indicators such as maximum yield and productivity and how to get these from a model parameters' search. In addition, the formulation of the corresponding optimization problem is presented. |
| 8 | Measurement data | Loading measurement data from spreadsheet files with subsequent creation of "Measurement" objects, focusing on three use cases that are based on: 1) Individual time vectors of varying lengths, with a shared time vector; 2) A shared time vector but missing data points for several measurements, and 3) Multi-replicate experiments with a shared time vector but missing data points. |
| 9 | Parallel parameter estimation (PPE) | Introduces PPE and the concept of continuation of an estimation job. |
| 10 | PPE – Optimizer comparison | Compares different optimization algorithms for PPE of a simple bioprocess model utilizing artificial noisy data. Comparison is based on runtimes and achieved losses for the given optimization problem. |
| 11 | PPE - Hyperparameter adjustment | Demonstrates how different parameter settings of the "de1220" and "compass_search" algorithms affect runtime and quality of the model calibration outcome. |
| 12 | PPE – Monte Carlo sampling | Introduces the application of PPE for Monte-Carlo sampling as method for non-linear error propagation. |
| 13 | PPE – Monte Carlo Sampling | In addition to the previous examples, the possibility to apply further constraints (beyond simple box bounds for parameters) is demonstrated. |
| 14 | Tracking specific rates during integration | Shows how specific rates can be derived and visualized as time-dependent performance indicators. |
| 15 | Non-negative states | For enforcing non-negative state values, events can be employed. Without, state values can take very small but negative numbers due to the operation mode of the integrator, which treats those values internally as zero (depending on the specified tolerances). |

## Notes

https://github.com/MicroPhen/pyFOOMB

### Competing Interest Statement

The authors have declared no competing interest.

